# *Notch3* is a genetic modifier of NODAL signalling for patterning asymmetry during mouse heart looping

**DOI:** 10.1101/2024.03.25.586543

**Authors:** Tobias Holm Bønnelykke, Marie-Amandine Chabry, Emeline Perthame, Audrey Desgrange, Sigolène M. Meilhac

## Abstract

The TGFβ secreted factor NODAL is a major left determinant required for the asymmetric morphogenesis of visceral organs, including the heart. Yet, when this signalling is absent, shape asymmetry, for example of the embryonic heart loop, is not fully abrogated, indicating that there are other factors regulating left-right patterning. Here, we used a tailored transcriptomic approach to screen for genes asymmetrically expressed in the field of heart progenitors. We thus identify *Notch3* as a novel left-enriched gene and validate, by quantitative in situ hybridization, its transient asymmetry in the lateral plate mesoderm and node crown, overlapping with *Nodal*. In mutant embryos, we analysed the regulatory hierarchy and demonstrate that *Nodal* in the lateral plate mesoderm amplifies *Notch3* asymmetric expression. The function of *Notch3* was uncovered in an allelic series of mutants. In single neonate mutants, we observe that *Notch3* is required with partial penetrance for the development of ventricles, in addition to its known role in coronary arteries. In compound mutants, we reveal that *Notch3* acts as a genetic modifier of *Nodal*, able to modulate heart looping direction and the curvature of the outflow tract. Whereas *Notch3* was previously associated with the CADASIL syndrome, its contribution to asymmetric organogenesis is now relevant to severe laterality defects such as the heterotaxy syndrome.

## Introduction

Left-right patterning of the embryo is essential for the formation of visceral organs (1,2). Anomalies in this process lead to the heterotaxy syndrome, a severe condition including complex congenital heart defects which determine the prognosis of patients (3). The mechanisms of symmetry-breaking are now well established in the mouse (4). Chirality of tubulin underlies the rotational movement of motile cilia in the pit of the left-right organiser, also referred to as the node. This generates a leftward fluid flow sensed by crown cells, resulting in the asymmetric expression of the left determinant NODAL in the lateral plate mesoderm. Genetic alterations to the formation of the node, ciliogenesis or NODAL signalling are associated with heterotaxy in both mice (2,5–8) and humans (9,10). NODAL signalling is then sensed by organ precursors to modulate morphogenesis, as shown in the intestine (11) and heart (12). In the absence of *Nodal*, some features are symmetrical (spleen, bronchi, lung lobes, atrial appendages), but many others (e.g heart looping, stomach, gut) retain some level of asymmetry, although abnormal (6,12–15). This indicates that *Nodal* is not always required to initiate asymmetry and that there are other factors of asymmetry in addition to NODAL signalling.

The heart is the first organ to undergo asymmetric morphogenesis in the embryo. This manifests as a rightward looping of the heart tube primordium, at E8.5 in the mouse (16). We have previously reconstructed the dynamics of this process and defined morphological and quantitative staging criteria from E8.5c to E8.5j, including the repositioning of ventricles from cranio-caudal to left-right, and the left displacement of the venous pole (17). We have shown that heart looping is primarily a buckling mechanism, when the heart tube elongates between fixed poles (17). We found that *Nodal* is not required for buckling and thus to initiate looping. In contrast, *Nodal* biases buckling, by amplifying and coordinating asymmetries independently at the two poles of the heart tube (12). Thus, in the absence of *Nodal* in the lateral plate mesoderm, asymmetries are reduced and the helical shape of the looped heart tube is abnormal. Because two asymmetries, at the arterial and venous poles, have a randomised orientation, *Nodal* mutants display 4 classes of abnormal heart loop. We observed that NODAL signalling is transient, before the formation of the heart tube (E8.5c-d), in precursor cells of the heart tube poles. Cardiac precursor cells are progressively incorporated into the heart tube, as a main driver of its elongation. Patterning of the field of cardiac precursors has been studied in terms of molecular profiling (18) and fate (19). However, the left-right patterning of cardiac precursors has remained centered on the *Nodal* pathway.

Other players of asymmetry have been identified, including BMP (20–22), HH (23), NOTCH (24–26) and WNT (27–29) signalling. Few of these are reported to be asymmetric: *Wnt3* in the left side of the node (28), and BMP signalling in the right lateral plate mesoderm (21). They were all shown to be required for *Nodal* asymmetry. Thus, the factors which can provide asymmetry besides *Nodal* have remained enigmatic.

NOTCH signalling is an example of a pathway required both for left-right asymmetry and for heart morphogenesis (30), but playing multiple roles. Among the four paralogues, *Notch2* is expressed in the node, and *Notch1* around the node in the caudal epiblast and presomitic mesoderm (24,31). *Notch1;Notch2* double mutants disrupt heart looping asymmetry, in keeping with a role in the left-right organiser (25). NOTCH1 and the transcriptional NOTCH effector RBPJ are required in the heart field for cardiomyocyte differentiation (32–34). *Notch1* is later expressed in the endocardium, where it is important to regulate myocardial trabeculation (35) and valve formation (36,37). *Notch3* mutant mice are viable and fertile (38), but display defects in the maturation of the smooth muscle wall of coronary arteries (39). *Notch3* is a broad marker of pericytes and controls the formation of smooth muscles in non-cardiac arteries and intestinal lacteals (40–42). Gain-of-function of NOTCH3 in CADASIL syndrome is responsible for smooth muscle degeneration in the brain (40,43,44). *Notch3* is also involved in the regulation of quiescence and stemness of neural, skeletal muscle and mammary stem cells (45–48). Finally, *Notch4* is expressed together with *Notch1* in endothelial cells to regulate vascular remodelling (49). Thus *Notch1/2* but not *Notch3/4* have been previously associated with left-right patterning, at the level of the left-right organiser.

To uncover asymmetric factors in organ specific precursors, rather than the left-right organiser, recent transcriptomic screens were performed on micro-dissected tissues excluding the node (50,51). Whereas NODAL signalling is transient (12,52), these screens were based on pooled embryos, which limits their resolution. The asymmetric candidate genes that have been identified have not yet been analysed for functional significance.

Here, we focused on asymmetric expression in the field of heart progenitors, at the time of heart looping. We compared paired left/right samples in single embryos to avoid confounding individual or stage variations. Our screen and quantitative in situ validations reveal *Notch3* as a novel gene enriched in the left lateral plate mesoderm. *Notch3* is co-expressed with *Nodal*, although with a slight time-delay and partial spatial overlap. In mutants, we show that *Notch3* asymmetry is amplified by NODAL signalling. Single *Notch3* mutants do not develop heterotaxy, but show ventricle and coronary artery defects after birth. Analysis of compound mutants indicate that *Notch3* is a genetic modifier of NODAL signalling during heart looping. Overall, we identify not only a novel asymmetric gene, but also a novel player in the left-right patterning of the lateral plate mesoderm, relevant to congenital heart defects.

## Results

### Identification of *Notch3* as a novel asymmetric marker

To screen for left-right asymmetric gene expression in cardiac cells, aside the *Nodal* pathway, we used a transcriptomic approach in the mouse embryo. To reduce confounding factors, the approach was targeted in space and time and left/right comparisons were conducted on single rather than pooled embryos. We focused on the heart field, given that it is a tissue which can be bisected at the midline and is patterned by NODAL signalling (12). In contrast, the heart tube changes its position during looping (17) and has a left-right origin difficult to dissect (19). We have previously shown that NODAL signalling is transient in heart precursors (12), highlighting the importance of specifically staging embryos. We focused on the E8.5f stage, when we have detected morphological asymmetry at the arterial pole of the heart tube (17). Thus, we micro-dissected the heart field of E8.5f wild-type embryos (Fig. 1A and S1A-D), in the dorsal pericardial wall and down to the second somite, in agreement with landmarks of fate mapping experiments (19) and profiling of single cardiac cells (18). Bulk RNA sequencing of paired left and right samples were compared to identify differential asymmetric expression in the heart field. As expected, targets of NODAL signalling, *Pitx2* and *Lefty2*, were significantly enriched on the left side (Fig. 1B). A total set of 597 genes were predicted to be differentially expressed (Table S1). Ingenuity Pathway analysis showed NOTCH signalling as the strongest asymmetric pathway and left-sided (Fig. 1C). We controlled the transcriptomic levels of individual genes associated with NOTCH in this differential analysis and their cardiac expression in a published single cell transcriptomic dataset at the same stage (18): whereas *Dtx4* was found to be expressed only in ectodermal cells (Fig. S1E), *Notch3*, *Hes1*, *Jag1*, *Ncstn* were identified as asymmetric candidate genes in the heart field (Fig. 1D).

**Fig 1.**
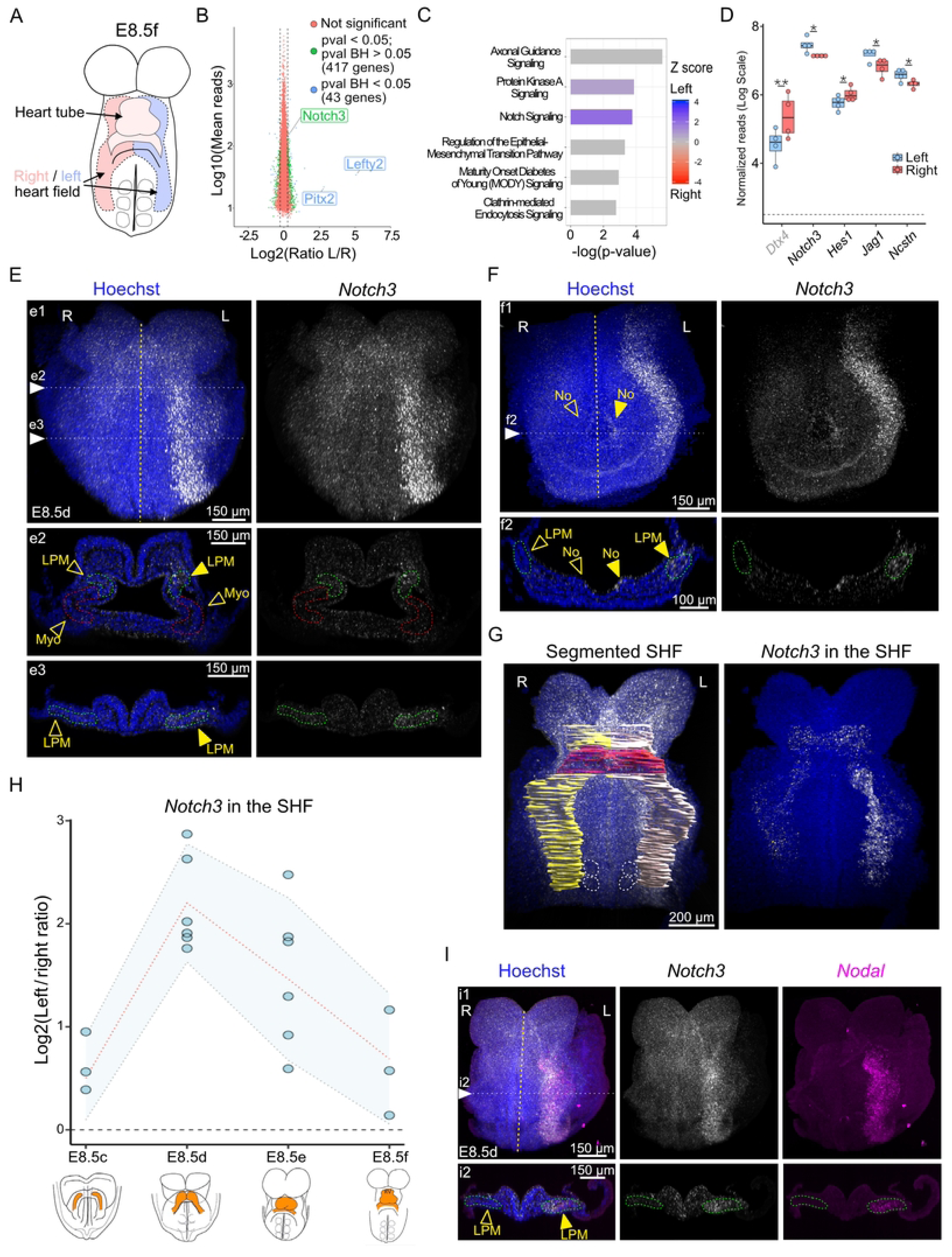
**RNA-sequencing of paired samples identifies a left-sided enrichment of *Notch3* at E8.5**. (A) Outline of the left (light blue) and right (light red) heart field micro-dissected for RNA-sequencing at the indicated stage of mouse heart looping. (B) MA-plot representing relative gene expression between paired left and right heart field at E8.5f. Non-significant differential expression is represented in red, differential expression in green (p-value<0.05), and blue (Benjamini-Hochberg (BH) corrected p-value<0.05, LimmaVoom, n=4 embryos). (C) Ingenuity Pathway Analysis on the gene list shown in Table S1 ordered according to significance and colour-coded for the activation state (z-score: blue, active pathway on the left; red, on the right). (D) Normalised read counts of genes involved in the *Notch* pathway in the left (blue) and right (red) heart field. The dotted line indicates the threshold of background expression. Whisker plots show the median, 25^th^– and 75^th^ quartiles (boxes), and the extreme data points (whiskers). *p-value < 0.05, ** Benjamini-Hochberg corrected p-value < 0.05 (LimmaVoom, n=4). (E-F) Expression of *Notch3* (white) detected by whole mount RNAscope ISH in E8.5d wild-type embryos, shown in frontal views (e1, cranial part of the embryo, f1, caudal part of the embryo) and transverse sections (e2-e3, f2, at the levels indicated in e1-f1) (n=6). Filled and empty arrowheads point to high and low expression, respectively. (G) Segmentation of the cardiac region in 3D images, to quantify gene expression in the left (white) or right (yellow) heart field. The heart tube is coloured in red. Somites are outlined by white dotted lines. Expression of *Notch3* within the segmented second heart field is extracted in the right panel. (H) Quantification of normalised *Notch3* asymmetric expression in the heart field at sequential stages. The means are shown on a red dotted line and standard deviations are in blue (n=3 at E8.5c, 6 at E8.5d, 6 at E8.5e, 3 at E8.5f). (I) *Notch3* (white) and *Nodal* (magenta) co-expression detected by double whole mount RNAscope ISH in E8.5d wild-type embryos, shown in a frontal view (i1) and transverse section (i2) (n=5). The midline is indicated by a yellow dotted line. HT, heart tube; L, left; LPM, lateral plate mesoderm (green dotted outline); Myo, myocardium (red 202 dotted outline); No, node; R, right; SHF, second heart field. See also Video S1.

We selected *Notch3* for further validation, because it is the gene with highest expression, it encodes a receptor and transcription co-factor central to NOTCH signalling. This *Notch* paralogue has not previously been associated with left-right patterning. We mapped *Notch3* expression in whole mount E8.5 wild-type embryos using sensitive RNAscope in situ hybridization. *Notch3* was detected throughout the lateral plate mesoderm, as well as in the crown of the node, with higher expression on the left side (Fig. 1E-F). *Notch3* was not expressed in the myocardium of the forming cardiac tube. We then assessed the dynamics of *Notch3* expression within the heart field (Fig. 1G). Our quantification shows that *Notch3* is transiently asymmetric, reaching 4.5-fold left-sided enrichment in the heart field at E8.5d, a stage when the myocardium bulges out to form a tube (Fig. 1H). Although *Nodal* asymmetry starts one stage earlier (E8.5c, (12)), *Notch3* and *Nodal* overlapped in the left lateral plate mesoderm at E8.5d (Fig. 1I, Video S1), and were found co-expressed in single cells (Fig. S1F). We have thus identified *Notch3* as a novel left-sided gene in the heart field.

### *Nodal* is required in the lateral plate mesoderm to amplify *Notch3* asymmetric expression

Given that *Notch3* asymmetry overlaps with that of *Nodal*, we investigated whether *Notch3* and *Nodal* depend on each other. NOTCH signalling has been shown to directly activate *Nodal* in the node, in *Dll1* and *Rbpj* mutants (26). In contrast, we found that *Nodal* was correctly patterned in the node and left lateral plate mesoderm of *Notch3^-/-^* mutants (Fig. S2A-B). In reverse, in mutants in which *Nodal* is inactivated in the lateral plate mesoderm but not in the node (12), *Notch3* expression was impacted (Fig. 2A-B). Quantification of *Notch3* expression at sequential stages in the heart field shows that *Notch3* asymmetry is severely decreased in *Nodal* mutants compared to controls (Fig. 2C). Yet, some significant asymmetry is still detectable. In keeping with a partial dependency of *Notch3* on *Nodal*, we observed that the expression domain of *Notch3* partially overlaps with that of *Nodal*, extending more medially and less laterally (Fig. S2C-D). Thus, *Nodal* is required in the left lateral plate mesoderm to amplify, rather than initiate, *Notch3* asymmetric expression.

**Fig 2.**
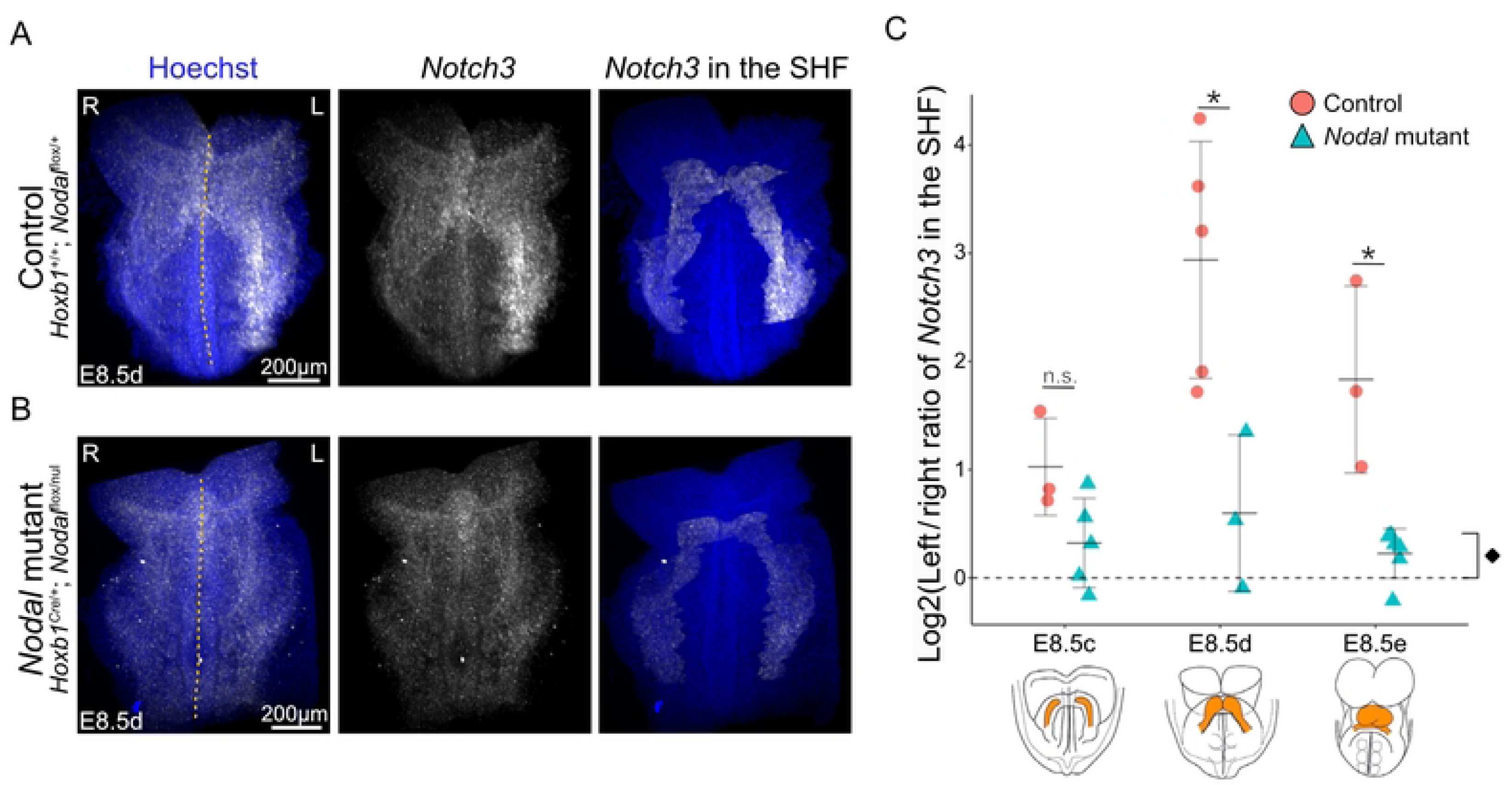
Decreased *Notch3* asymmetry in *Nodal* mutants. (A-B) Whole mount RNAscope ISH of *Notch3* in control (A) and *Nodal* mutants (B) at E8.5d in frontal views. Expression of *Notch3* within the segmented heart field is extracted in right panels. The midline of the embryo is indicated by a yellow dotted line. (C) Corresponding quantification of normalised *Notch3* asymmetric expression in the heart field at sequential stages in littermate controls (n= 3 at E8.5c, 5 at E8.5d, 3 at E8.5e) and *Nodal* mutants (n= 5 at E8.5c, 3 at E8.5d, 6 at E8.5e). Means and standard deviations are shown. *p-value<0.05 between controls and mutants, ♦p-value<0.05 to compare mutant levels with a symmetry hypothesis (Log2 ratio =0) (Pairwise Mann-Whitney Wilcoxon tests). L, left; R, right; SHF, second heart field.

### Single *Notch3* mutants have normal heart looping but later congenital heart defects

We next investigated the role of *Notch3* in heart development, in mutants inactivated for it (Fig. S3). Given its asymmetric expression when the cardiac tube forms, we looked for a potential effect on the rightward looping of the heart tube. E9.5 mutant embryos were collected at Mendelian ratio (Fig. 3A) and no heart looping defect was detected, both qualitatively (Fig. 3B-D) and quantitatively (Fig. 3E-F). *Notch3^-/-^* mutant embryos were indistinguishable from controls, with a normal orientation of the right ventricle-left ventricle axis relative to the midline, and a normal left displacement of the venous pole.

**Fig 3.**
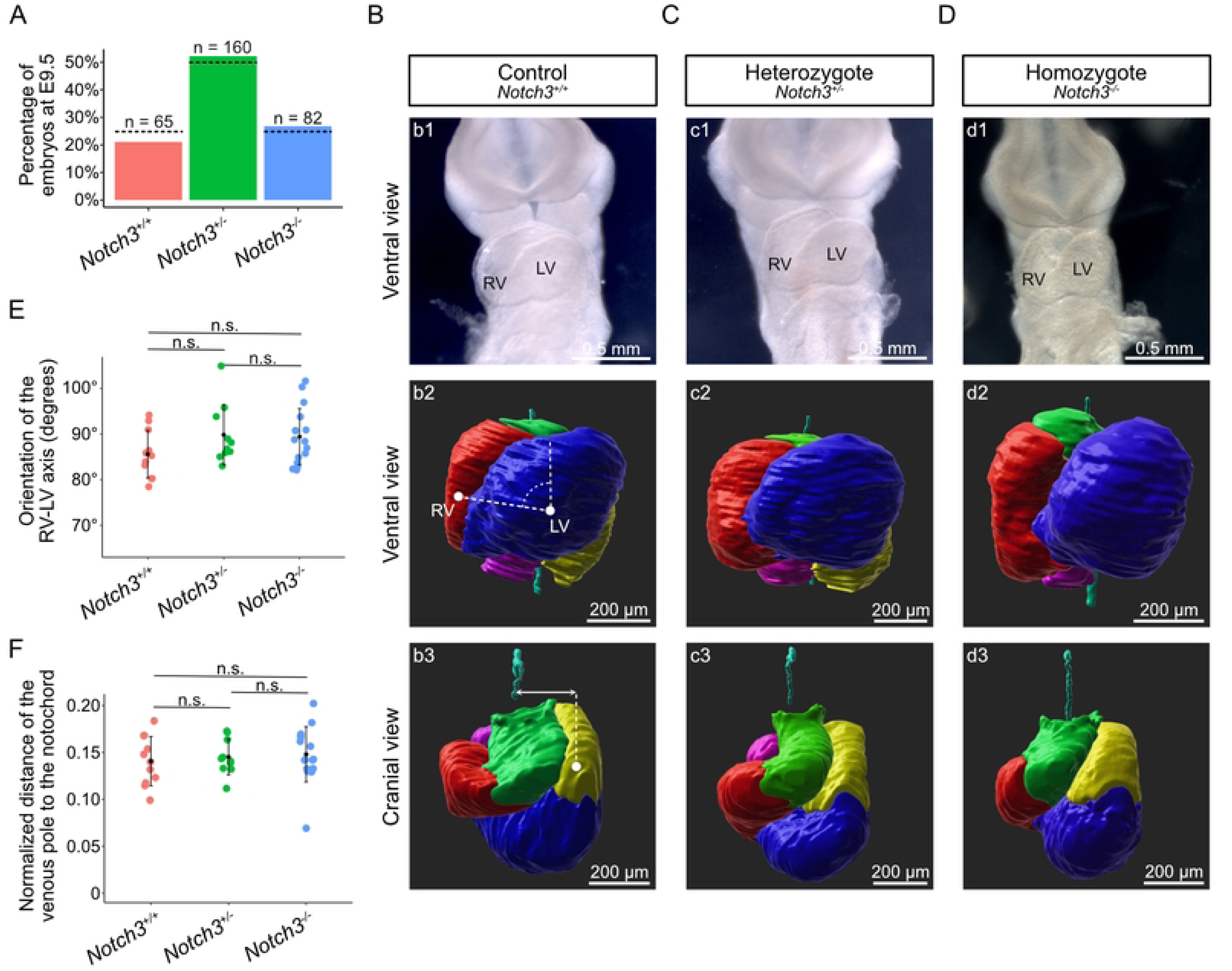
Normal heart looping in single *Notch3* mutants. (A) Histogram showing the percentage of genotypes recovered in E9.5 litters of *Notch3+/-* x *Notch3+/-* crosses. The observed frequency is not significantly different from the expected Mendelian ratio (dotted lines) (p-value = 0.30, chi-square test, n as indicated). (B-D) Brightfield ventral views (b1-d1), and 3D segmented heart tubes shown in ventral (b2-d2) and cranial (b3-d3) views, of littermate *Notch3+/+* (B), *Notch3+/-* (C) and *Notch3-/-* (D) E9.5 embryos. Cardiac regions are colour coded: outflow tract in green, right ventricle in red, left ventricle in blue, atrioventricular canal and left atrium in yellow, right atrium in magenta. The notochord (cyan) is used as a reference axis to align samples. (E) Quantification of the orientation of the RV/LV axis relative to the notochord, as schematised in b2. (F) Quantification of the displacement of the venous pole relative to the notochord, normalised by the tube length, as schematised in b3. Means and standard deviations are shown (Pairwise Mann-Whitney Wilcoxon tests with Benjamini-Hochberg correction, n=11 *Notch3+/+*, 10 *Notch3+/-* and 16 *Notch3-/-*). LV, left ventricle; n.s., non-significant; RV, right ventricle.

We then collected *Notch3* mutants after birth. In agreement with previous reports (38), they were collected at Mendelian ratio (Fig. 4A). *Notch3^-/-^* mutants did not show heterotaxy, unlike *Nodal* mutants, but had congenital heart defects with partial penetrance (67%), including peri-membranous and muscular ventricular septal defects, thinning of the right ventricular wall and coronary artery dilatation (Fig. 4B-C). Defects of coronary arteries are in keeping with fetal expression of *Notch3* in pericytes and smooth muscle cells (Fig. S4). Overall, our results show that *Notch3* is required for ventricle and coronary artery development. However, removal of *Notch3* alone does not uncover a role in left-right asymmetric organogenesis.

**Fig 4.**
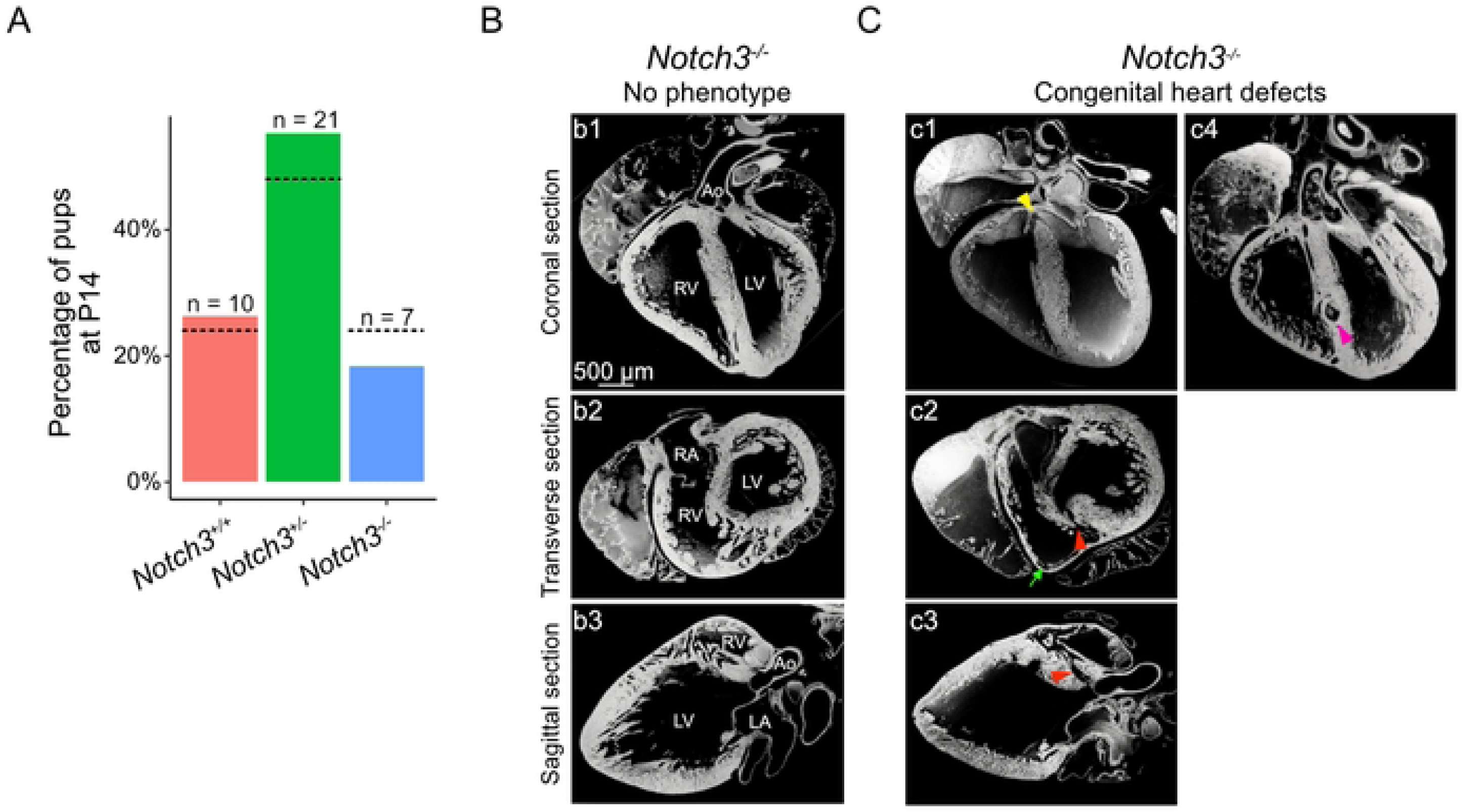
Congenital heart defects in *Notch3* mutants. (A) Histogram showing the percentage of genotypes recovered in P14 litters of *Notch3+/-* x *Notch3+/-* crosses. The observed frequency is not significantly different from the expected Mendelian ratio (dotted lines) (p-value = 0.64, chi-square tests, n as indicated). (B-C) Sections of *Notch3-/-* mutant hearts at P0, imaged by HREM. An example of a mutant heart with no phenotype is shown (B, n=4/12). Defects (C) include peri-membranous ventricular septal defect (VSD) (yellow arrowhead, n=1/12), muscular VSD (red arrowheads, n=4/12), dilatation of the septal coronary artery (pink arrowhead, n=1/12) and thin RV wall (green arrow, n=3/12). AVC, atrioventricular canal; Ao, aorta; LA, left atrium; LV, left ventricle; RA, right atrium; RV, right ventricle.

### *Notch3* is a genetic modifier of *Nodal* in heart looping

Since *Notch3* overlaps with and is amplified by *Nodal*, we investigated their interaction in the context of left-right asymmetry. We generated double heterozygote mutants and quantified heart looping. Compared to single heterozygotes (Fig. 5A-B), *Notch3^+/-^; Hoxb1^Cre/+^; Nodal^flox/+^* double heterozygotes (Fig. 5C) did not show any heart looping defects, including normal orientation of the right ventricle-left ventricle axis relative to the midline, and normal left displacement of the venous pole (Fig. 5D-E). Thus, we did not detect a genetic interaction between *Notch3* and *Nodal*.

**Fig 5.**
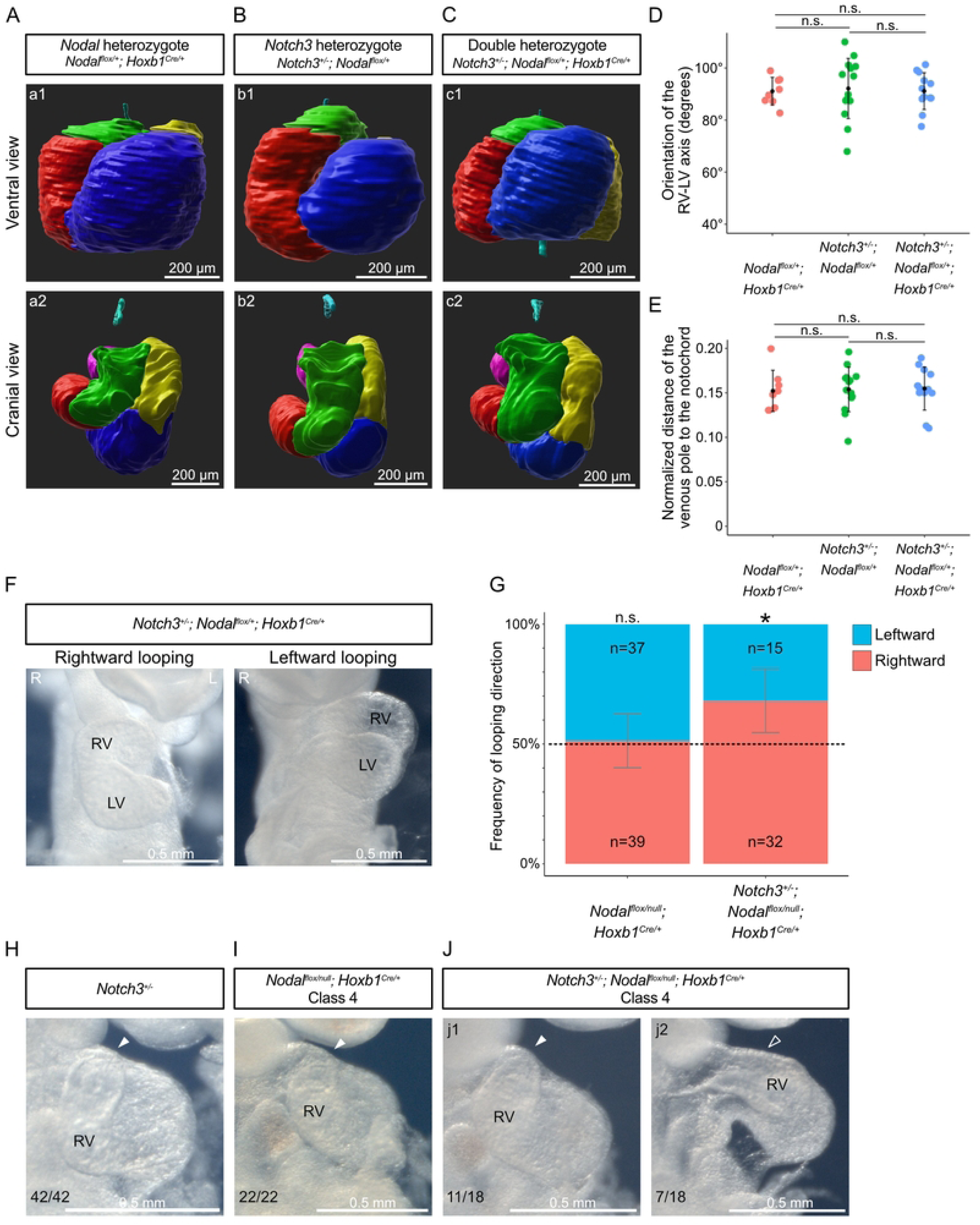
Heart looping variations in *Notch3* and *Nodal* compound mutants. (A-C) 3D segmented heart tubes shown in ventral (a1-c1) and cranial views (a2-c2) of single *Nodal* heterozygote (A), single *Notch3* heterozygote (B) and double heterozygote (C) embryos at E9.5. Cardiac regions are colour coded as in Fig. 3. The notochord (cyan) is used as a reference axis to align samples. **(D)** Quantification of the orientation of the RV/LV axis relative to the notochord. **(E)** Quantification of the displacement of the venous pole relative to the notochord, normalised by the tube length. Means and standard deviations are shown. (Pairwise Mann-Whitney Wilcoxon tests with Benjamini-Hochberg correction, n=8 *Hoxb1Cre/+; Nodalflox/+,* 14 *Notch3+/-; Nodalflox/+,* 12 *Notch3+/-; Hoxb1Cre/+; Nodalflox/+*). **(F)** Brightfield images of homozygote *Nodal* mutants with decreased *Notch3* (*Notch3+/-; Hoxb1Cre/+; Nodalflox/null*) at E9.5, in a ventral view. **(G)** Corresponding quantification of their looping direction frequency (right bar), which, in contrast to *Nodal* mutants with normal *Notch3* levels (left bar), differs from a randomised looping direc[on (dotted line). *p-value<0.05 (Chi-square test with Yates’ continuity correction, n=76 *Hoxb1Cre/+; Nodalflox/null* and 47 *Notch3+/-; Hoxb1Cre/+; Nodalflox/null*). **(H-J)** Comparison of Class 4 abnormal heart loops in homozygote *Nodal* (I), *Nodal; Notch3* compound (J) mutants compared to control *Notch3+/-* embryos (H), seen in brightfield right-sided views. Filled and empty arrowheads point to curved and straight outflow tract, respectively. L, Left; LV, left ventricle; R, Right; RV, right ventricle.

We then analysed the role of *Notch3* in a sensitized background. We generated a large cohort of mutants, with a knock-down of *Notch3* within the context of *Nodal* inactivation in the left lateral plate mesoderm. *Notch3^+/-^; Hoxb1^Cre/+^; Nodal^flox/null^* mutants lost the randomized direction of heart looping, typical of *Nodal* mutants (Fig. 5F-G). Rightward looping was more frequent when *Notch3* was reduced in *Nodal* mutants. However, this did not correspond to a rescue, since all rightward mutant heart loops were abnormal, compared to controls. Compared to *Nodal* mutants, additional defects in the outflow tract were observed with partial penetrance. The outflow tract was straighter in 40% (7/18) of *Notch3^+/-^; Hoxb1^Cre/+^; Nodal^flox/null^* mutants within looping Class 4 (Fig. 5H-J), which is the less severe class. These genetic analyses thus uncover a role for *Notch3* as a genetic modifier of *Nodal* during left-right asymmetric organogenesis.

## Discussion

We have identified *Notch3* as a novel left asymmetric factor. We show that it is amplified by NODAL signalling. We do not detect a genetic interaction between *Nodal* and *Notch3*, but rather show that *Notch3* acts as a genetic modifier of *Nodal*. Knock-down of *Notch3* exacerbates looping defects observed in *Nodal* mutants. Single *Notch3* mutant neonates have other cardiac defects in coronary arteries, transmural growth and septation of ventricles.

Left-right asymmetry is mainly analysed in terms of patterning of the major left determinant, *Nodal*, and its downstream pathway including *Lefty2* and *Pitx2* (4,53,54). However, in the absence of *Nodal*, organ asymmetry is not fully abrogated (6,13–15). Heart looping, which is the first asymmetric morphogenesis, still occurs to some extent in *Nodal* mutants, producing helical heart tubes, even if abnormal (12). This indicates that there are other factors than NODAL signalling required for heart asymmetry. BMP (20–22), HH (23), NOTCH (24–26) and WNT (27–29) signalling have been shown to be required upstream of *Nodal* for asymmetry. Transcriptomic screens have been performed to identify asymmetric genes in embryos outside the node or in cardiac precursors (50,51). Yet, asymmetric candidate genes were either not validated or not analysed functionally, leaving open the question of which are genuine asymmetry factors. Here, we have used a transcriptomic screen tailored spatially in the heart field, temporally at the stage of the first morphological asymmetry, and specifically on left-right asymmetry based on paired comparison in single embryos. This has successfully identified a novel left factor, *Notch3*. We have characterised by quantitative and whole mount sensitive in situ hybridisation its dynamic spatio-temporal expression pattern relative to *Nodal*. And we have demonstrated its role in left-right asymmetry based on an allelic series of mutants. Similarly to *Nodal*, *Notch3* is transiently asymmetric, highlighting the importance of careful staging to monitor left-right patterning. In contrast to *Nodal*, *Lefty2* and *Pitx2,* which are exclusively expressed on the left of the lateral plate mesoderm, *Notch3* is also expressed on the right, with a 4.5-fold enrichment on the left side. Thus, quantitative and sensitive approaches are critical to uncover factors which have lower asymmetry levels compared to the *Nodal* pathway.

*Nodal* plays multiple roles in asymmetry. *Nodal* is well known to act as a bias, able to set the laterality of asymmetry, so that its absence leads to a randomized orientation. This is the case for the direction of heart looping (12) or the stomach position (14). In addition, *Nodal,* upstream of *Pitx2*, acts as a regulator of identity, able to confer a left anatomical structure to the atria and lungs or to induce the formation of the spleen (14,55,56). Finally, we had shown that *Nodal* can also act as an amplifier of asymmetry (12). This had been detected at a morphological level, with a reduced rotation of the arterial pole during heart looping in *Nodal* mutants. We now show this at a molecular level, with the amplification of *Notch3* asymmetry by *Nodal*.

NOTCH signalling is required at different levels for left-right asymmetry. *Notch1/2* were shown previously to play a role in the formation of the left-right organiser and the induction of *Nodal* (24–26). We now identify the role of another NOTCH paralogue, NOTCH3. Whereas *Notch1* is expressed around the node in the caudal epiblast and presomitic mesoderm, and *Notch2* in the node pit (24,31), we have detected *Notch3* in the node crown and lateral plate mesoderm. Our conditional mutants permit to dissect a role of *Notch3* in the lateral plate mesoderm for left-right asymmetry. How *Notch3* regulates heart looping direction and the curvature of the outflow tract remains an open question. *Notch3* regulates muscle differentiation and maturation in other tissues, such as the smooth muscles of arteries, coronary arteries and intestinal lacteals, or skeletal muscles, and is expressed in precursors cells such as pericytes or satellite stem cells (39–42,45). It will be interesting to investigate whether *Notch3* in the heart field similarly regulates the differentiation and maturation of the cardiac muscle to impact heart looping. Elongation of the heart tube is important for heart looping, as a driver of a buckling mechanism (17). Thus, asymmetries in myocardial cell differentiation, when precursor cells incorporate into the heart tube, can provide a left-right bias to orient looping. We have shown previously that *Nodal* modulates a large set of genes involved in myocardial cell differentiation, some of which like *Tnnt1* are asymmetrically expressed within the wild-type heart tube (12). Whether *Notch3* acts similarly remains to be investigated. For later transmural growth and septation of ventricles, it is possible that a potential role of *Notch3* on muscle differentiation and maturation intervenes. However, since *Notch3* is expressed in cardiac precursors and not cardiomyocytes, that would imply a long delay from expression to phenotype. Another possibility is that ventricle growth in *Notch3* mutants is a secondary consequence of defective coronary vascularisation (57).

Although *Notch3* expression is sensitive to *Nodal*, our observations support the idea that *Notch3* is not a component of the *Nodal* pathway. The expression pattern of *Notch3* differs from *Nodal*, with a right sided and more medial left sided localization. In contrast to *Pitx2* or *Lefty2* (12), *Notch3* expression is not erased in *Nodal* mutants. Functionally, we do not detect a genetic interaction between *Notch3* and *Nodal*, and show that knock-down of *Notch3* can exacerbate the phenotype of *Nodal* mutants. Thus, *Notch3* comes out as a novel genetic modifier of *Nodal*. The underlying mechanism is still unknown. A link between the two pathways has been detected based on the direct binding of NOTCH3 to the NODAL co-receptor TDGF1, which enhances NOTCH proteolytic maturation and sensitization to ligand-induced NOTCH signalling (58). Given that the disease associated with *Nodal* dysfunction, heterotaxy, has a heterogenous phenotypic spectrum, the identification of genetic modifiers of known genes involved in heterotaxy opens novel perspectives to fill gaps in the genetic diagnosis of 60% of patients (59).

## Methods

### EXPERIMENTAL MODEL AND SUBJECT DETAILS

C56Bl6J mice were used as wild-type embryos. *Notch3^tm1Grid/tm1Grid^* mutants (abbreviated *Notch3^-/-^*, (38)) were kept in a C56Bl6J background; they are viable and fertile as homozygotes. *Nodal^null/+^; Hoxb1^Cre/+^* males (12) were maintained in a mixed genetic background and crossed with *Nodal^flox/flox^* females (60) to generate *Nodal* conditional mutants. Both male and female embryos were collected and used randomly for experiments, except for RNA sequencing, in which only male embryos were used to reduce variability. Embryonic day (E) 0.5 was defined as noon on the day of vaginal plug detection. Embryonic stages were determined based on the morphology of the heart according to (17) and (12). Somite numbers were evaluated from brightfield images; samples with less than 18 somites were excluded from heart looping quantifications. All embryos were genotyped by PCR, using primers listed in Table S2. Animals were housed in individually ventilated cages containing tube shelters and nesting material, maintained at 21°C and 50% humidity, under a 12h light/dark cycle, with food and water ad libitum, in the Laboratory of Animal Experimentation and Transgenesis of the SFR Necker, Imagine Campus, Paris. Animal procedures were approved by the ethical committees of the Institut Pasteur, Université Paris Cité and the French Ministry of Research.

### METHOD DETAILS

#### RNA isolation and sequencing

Paired left and right heart fields of E8.5f embryos were micro-dissected, from below the headfolds to the end of the second somite (19), after removal of the heart and the back. The tissue was flash frozen in liquid nitrogen. All samples were collected within 1h30 of sacrificing the mother. Total RNA was extracted in TRIzol-Chloroform and purified using the RNeasy micro kit (QIAGEN) including DNAse treatment. RNA quality and quantity were measured on Fragment Analyzer (Agilent Technologies). All RQN were >9.7. The libraries were established using the Nugen Universal Plus mRNA-Seq kit, using 15ng of total RNA per sample. The oriented cDNAs produced from the poly-A+ fraction were PCR amplified (15-18 cycles). An equimolar pool of the final indexed RNA-Seq libraries was sequenced on an Illumina NovaSeq6000, with paired-end reads of 130 bases and a mean sequencing depth of 37.15 million reads per sample.

#### Embryo dissection

Embryos were dissected in 1xDPBS and fixed in 4% paraformaldehyde either for 6h at room temperature or 24h at 4°C. Yolk sac or tail pieces were collected for genotyping. For embryos dissected at E9.5, hearts were arrested in diastole by treatment with cold 250mM KCl for 5 minutes. Fixed embryos were gradually dehydrated into methanol and stored at –20°C. For pups collected at P0, they were euthanised, immerged in a cardioplegia solution (110mM NaCl, 16mM KCl,16mM MgCl2, 1.5mM Cacl2, 10mM NaHCO3) for 5 minutes and fixed in 4% PFA 24h at 4°C.

#### RNA in situ hybridization (ISH)

ISH was performed whole mount as in (12). *Wnt11* and *Bmp2* antisense riboprobes were transcribed from plasmids. Signals were detected by alkaline phosphatase (AP)-conjugated anti-DIG antibodies (1/2000), which were revealed with the BM purple (magenta) substrate. The samples were washed in 1x DPBS, post-fixed and imaged by HREM.

Wholemount RNAscope ISH were performed using the Mutliplex Fluorescent v2 Assay (Biotechne) and the protocol of (61). *Notch3* (425171-C1, 425171-C2) and *Nodal* (436321-C1) probes were used. Amplification steps were performed using the TSA cyanine5 and cyanine3 amplification kit. Hoechst (1/1000) was used as a nuclear counterstain. Samples were then transferred in R2 CUBIC clearing reagents and embedded in R2 reagent containing agarose (62). Multi-channel 16-bit images were acquired with a Z.1 lightsheet microscope and a 20X/1.0 objective.

#### RT-qPCR

Reverse transcription was performed on RNA isolated from entire (left and right) micro-dissected heart fields using a Reverse Transcription kit (QuantiTect, Qiagen). Quantitative PCR was carried out using the ViiA7 real-time PCR system. mRNA expression levels were measured relatively to *Polr2b* and normalised with a reference cDNA sample, taken as a pool of 7 whole embryos at stage E8.5c-g, using the standard ΔΔCt method. Primers are listed in Table S2.

#### High Resolution Episcopic Microscopy (HREM)

E9.5 embryos or P0 hearts were imaged in 3D by HREM after embedding in methacrylate resin (JB4) containing eosin and acridine orange as contrast agents (12). Two channel images of the surface of the resin block were acquired using the optical high-resolution episcopic microscope and a 1X Apo objective repeatedly after removal of on average 1.75 µm or 2.63 µm thick sections. The datasets comprise 742-1869 images of 0.90-4.48 µm resolution in x and y depending on the stage. Icy (63) and Fiji (ImageJ) softwares were used to crop and scale the datasets. 3D reconstructions and analysis were performed using Imaris.

### QUANTIFICATION AND STATISTICAL ANALYSIS

#### Bioinformatics Analyses of bulk RNA sequences

FASTQ files were mapped to the ENSEMBL Mouse GRCm38/mm10 reference using Hisat2 and counted by featureCounts from the Subread R package. Due to a high number of duplicates (on average 89%), duplicates were excluded. Flags were computed from counts normalized to the mean coverage. All normalized counts < 20 were considered as background (flag 0) and ≥ 20 as signal (flag = 1). P50 lists used for the statistical analysis gather genes showing flag=1 for at least half of the samples. For the analysis of gene expression, normalized read counts below 250 were considered as background, based on known markers included or excluded from the micro-dissected tissue (Fig. S1B-D). Differential gene expression analysis was performed using three independent methods (DESeq2, edgeR and LimmaVoom) followed by thresholds on absolute fold change ≥1.2 and p-value ≤0.05, leading to 597 genes (Table S1). Differential gene expression with LimmaVoom is plotted, and grouped in three categories depending on p-value and corrected p-value (Benjamini-Hochberg procedure). For each differential analysis method, functional analyses were carried out using the Ingenuity Pathway analysis on the list of differentially expressed genes, using delta between conditions and p-value of differential analysis. Transcriptomic data have been deposited in NCBI Gene Expression Omnibus (GEO) with the accession number GSE237126.

#### Bioinformatics Analyses of published single cell RNA sequences

Data from single cardiac wild-type cells at E8.5 (18) were downloaded from https://crukci.shinyapps.io/heartAtlas/ analysed using scran (64) and visualized using Seurat v3.1.2 (65). Data from single cardiac wild-type cells at E11.5-P9 (66) were downloaded from https://www.ncbi.nlm.nih.gov/geo/query/acc.cgi?acc=GSE193346, and visualized similarly.

#### Quantification of RNAscope ISH signal

3D images were analyzed using Imaris. For quantification of left-right asymmetry in the second heart field, the second heart field was manually segmented in the lateral plate mesoderm layer, using headfolds and the second somite pair as cranial and caudal boundaries, respectively, and separated into left and right following the midline (using neural tube closure as a landmark). Both Hoechst signal and the signal of the gene of interest were extracted. To account for potential biases between the left and right during imaging, the spot detector function was used on the Hoechst channel to generate a normalisation score between the two sides, using as a threshold a coverage of 50% of spots on each side. To calculate the left/right ratio of the gene of interest, its number of spots were measured on the left and right sides, normalised to Hoechst signal and to the volume of the segmented region.

#### Quantification of the geometry of the heart tube at E9.5

HREM images were used to segment the different compartments of the heart tube in Imaris. Eight landmarks along the tube were extracted and used for quantifications as described previously (12): one at the exit of the outflow tract, one at the sulcus between the two ventricles, and one at the bifurcation of the two atria (taken as the venous pole). The five other landmarks are centroids of cardiac regions: outflow tract, right ventricle, left ventricle, atrioventricular canal and left atrium, and right atrium, segmented according to the expression patterns of *Wnt11* and *Bmp2* and anatomical landmarks such as cushion boundaries or the interventricular or interatrial sulcus. Heart shapes were aligned in 3D using an in-house MATLAB code so that the Z-axis corresponds to the notochord axis and the x-axis to a perpendicular dorso-ventral axis. The orientation of the axis between the left and right ventricles, the distance of the venous pole to the notochord and the tube length were measured as in (17).

#### Phenotyping of congenital heart defects

Hearts of *Notch3* mouse mutants were phenotyped at P0 in 3D images acquired by HREM, based on the segmental approach (67) and IPCCC ICD-11 clinical code.

#### Statistical Analyses

Statistical tests and p-values are described in figure legends and Table S3. Group allocation was based on PCR genotyping. All sample numbers (n) indicated in the text refer to biological replicates, i.e. different embryos or different cells. Investigators were blinded to allocation during imaging and phenotypic analysis, but not during quantifications. Pairwise Mann-Whitney-Wilcoxon tests were used to compare experimental groups, with a Benjamini-Hochberg correction applied on p-values when more than 2 populations are compared. A chi-square test was used to compare percentage distributions, or a chi-square test with Yates’ continuity correction for comparison with an expected value. Tests were performed with R and Excel. Embryos which had been damaged during experiments were excluded for quantification.

## Abbreviations

Ao: Aorta
AVC: Atrioventricular canal
BH: Benjamini-Hochberg
BMP: bone morphogenetic protein
E: Embryonic day
HREM: High Resolution Episcopic Microscopy
HT: Heart tube
ISH: In situ hybridisation
IPCCC ICD: International Paediatric and Congenital Cardiac Code and International Classification of Diseases
L: Left
LA: Left atrium
LPM: Lateral plate mesoderm
LV: Left ventricle
Myo: Myocardium
No: Node
n.s.: non-significant
NT: Neural tube
P: Postnatal day
R: Right
RA: Right atrium
RT-qPCR: Reverse transcription quantitative polymerase chain reaction
RV: Right ventricle
SHF: Second Heart Field
TGFβ: Transforming growth factor-β
VSD: Ventricular Septal Defect

## Acknowledgements

We thank V. Benhamo, L. Guillemot, M. Bertrand, M. Cavaignac, M. Franco, J. Terret, N. Agueeff for generous technical assistance; L. Bally-Cuif, H. Marlow, P. Seymour, N. Kurpios and S. Fre for insightful discussions; A. Joutel for providing *Notch3^tm1Grid^* mice; D. Conrozet, S. Ameur and the histology platform of the SFR Necker; the cell imaging platform; C. Bole-Feysot, M. Parisot and M. Zarhrate of the genomics platform; N. Cagnard of the bioinformatics platform; the LEAT animal facility and H. Varet of the Pasteur Bioinformatics and Biostatistics Hub.

## Author Contributions

Conceptualization, S.M.M., A.D., T.H.B.; Formal Analysis, T.H.B., A.D., M-A.C., E.P.; Funding Acquisition, S.M.M.; Investigation, T.H.B., A.D., M-A.C.; Methodology, T.H.B., A.D., E.P., S.M.M.; Project Administration, S.M.M.; Software, T.H.B., E.P.; Supervision, S.M.M., A.D., T.H.B.; Visualization T.H.B., A.D., M-A.C., E.P.; Writing – Original Draft, S.M.M.; Writing – Review & Editing, all authors;

## Additional information

**Funding**. This work was supported by core funding from the Institut Pasteur and INSERM, state funding from the Agence Nationale de la Recherche under ‘‘Investissements d’avenir’’ program (ANR-10-IAHU-01, ANR-10-LABX-73-01 REVIVE), the Philanthropy Department of Mutuelles AXA through the Head and Heart Chair, a grant from the ANR (ANR-18-CE15-0025-03) and the MSD-Avenir fund (Devo-Decode project) to S.M.M. T.H.B. was supported by the Pasteur – Paris University (PPU) International PhD Program and the FRM, M-A.C. is a student from the FIRE PhD program funded by the Bettencourt Schueller foundation and the EURIP graduate program (ANR-17-EURE-0012). A.D. is an INSERM researcher and S.M.M. an INSERM research director.

## Supporting Information

**Figure S1 related to Fig. 1. Transcriptomic approach. (A)** Brightfield images of wild-type embryos used for RNA sequencing at E8.5f. In the left panel, an outline of the dissected areas is shown. The identification number of embryos is given. **(B)** Normalised read counts of genes used to validate the threshold of expression in the transcriptomic analysis. The osteocyte gene *Dmp1*, inner ear marker *Oc90*, neuronal marker *Neurod1* are used as negative controls and *Mmp9* as a positive control, lowly expressed in left heart progenitors. Whisker plots show the median, 25^th^– and 75^th^ quartiles (boxes), and the extreme data points (whiskers). **(C)** Normalised read counts of genes used as markers to control sample micro-dissection. *En2, Wnt8b* are anterior markers, *Hoxa5, Hoxb6* posterior markers, *Rfx4*, *Tfap2b* back markers, *Isl1, Six2, Fgf8* second heart field markers, *Tbx5, Mab21l2* juxta-cardiac field (JCF) markers and *Myh7b, Myoz1, Tnnt1* cardiomyocyte markers. The dotted line indicates the threshold of background expression. **(D)** Normalised read counts of genes used as markers to validate the left-right dissection of samples. NODAL targets *Lefty2* and *Pitx2,* as well as *Six2* label the left side. *Nodal* is turned off at E8.5f. *p-value between the left and right sides < 0.05, *** Benjamini-Hochberg corrected p-value < 0.00001 (LimmaVoom, n=4). **(E)** Violin plot of *Dtx4* expression in single cells at E8.5 from (18), clustered as annotated (n= 89 Ec2, 59 Me2, 713 Me3, 221 Me4, 355 Me5, 65 Me6, 514 Me7). Dots are normalised reads per cell. **(F)** Violin plot of *Notch3* expression in single *Nodal*-negative (n=1559) and *Nodal*-positive (n=309) cells of cardiac clusters (Me3-7) of (18) at E8.5 (stage 1 to Late Heart Tube, wild-type embryos). 75% (234/309) of *Nodal*-positive cells also express *Notch3*.

**Figure S2 related to Fig. 2. Normal *Nodal* expression in *Notch3* mutants. (A-B)** Whole mount RNAscope ISH of *Nodal* in *Notch3^-/-^* mutant embryos at E8.5c in anterior (A) and posterior (B) frontal views. The midline of the embryo is indicated by a yellow dotted line. A transverse section at the level of the node (see b1) is shown in b2. Filled and empty arrowheads point to high and absent expression, respectively. **(C-D)** Transverse section of the left lateral plate mesoderm, labelled by double wholemount RNAscope ISH of *Nodal* (red) and *Notch3* (white). The region of *Notch3* and *Nodal* co-expression is outlined in yellow (n=3). L, left; LPM, lateral plate mesoderm; No, node; NT, neural tube; R, right.

**Figure S3 related to Fig. 3. *Notch3* inactivation in mutants. (A)** Schema of *Notch3* alleles in wild-types and *Notch3^-/-^* mutants, indicating the localisation of exons and primers used for RT-qPCR. **(B)** Relative expression of *Notch3* in micro-dissected heart fields of littermate wild-types (n=5 E8.5e, 3 E8.5g), *Notch3^+/-^* (n=6 E8.5e, 6 E8.5g) and *Notch3^-/-^* (n=6 E8.5e, 4 E8.5g) embryos, quantified by RT-qPCR using the indicated primer pairs and normalised to wild-types. **p-value<0.01, ***p-value<0.001 (Pairwise Mann-Whitney Wilcoxon tests with Benjamini-Hochberg correction).

**Figure S4 related to Fig. 4. *Notch3* expression after heart looping.** Violin plot of *Notch3* expression in single cardiac cell transcriptomic between E11.5 and P9 (from (66)), clustered as annotated (n= 5422 atrial cardiomyocytes, 10493 ventricular cardiomyocytes, 2361 endocardium, 1176 vascular endocardium, 1012 epicardium, 3309 fibroblast like cells, 182 smooth muscle cells, 349 pericytes, 1032 macrophages, 33 B cells, 42 T cells, 17 dendritic cells, 23 natural killer cells, 22 neutrophils, 244 blood cells). Dots are normalised reads per cell.

**Video S1 related to Fig. 1. Co-expression of *Notch3* and *Nodal*.** *Notch3* (white) and *Nodal* (magenta) double whole-mount RNAscope ISH in a wild-type E8.5d embryo imaged in 3D by lightsheet microscopy.

**Table S1 related to Fig. 1.** List of 597 differentially expressed genes used for Ingenuity Pathway analysis, based on a fold change ≥1.2 and p-val ≤0.05 with either the LimmaVoom, DESeq2 or edgeR methods. Flags indicate the number of samples in which normalised read counts are ≥ 20 (n=4 embryos). Lf, left heart field at E8.5f; Rf, right heart field at E8.5f.

**Table S2. List of primers** used for genotyping and RT-qPCR.

**Table S3. Data.** Numerical data used to generate figure graphs along with statistical tests.

## Notes

### Competing Interest Statement

The authors have declared no competing interest.

